# H7N6 Low Pathogenic Avian Influenza outbreak in commercial turkey farms in Chile caused by a native South American Lineage

**DOI:** 10.1101/527069

**Authors:** C. Mathieu, A. Gonzalez, A. Garcia, M. Johow, C. Badia, C. Jara, P. Nuñez, V. Neira, N.A. Montiel, M.L. Killian, B.P. Brito

**Affiliations:** Servicio Agrícola y Ganadero (SAG), Laboratorio y Estación Cuarentenaria de Lo Aguirre, Santiago, Chile; Departamento de Medicina Preventiva, Facultad de Ciencias Veterinarias y Pecuarias, Universidad de Chile, La Pintana, Santiago, Chile; National Veterinary Services Laboratories, Science, Technology and Analysis Services, Veterinary Services, Animal and Plant Health Inspection Service, U.S. Department of Agriculture, Ames, Iowa, USA; The ithree institute, University of Technology Sydney, PO Box 123, Broadway, NSW 2077, Australia

**Keywords:** Influenza in birds, Disease Outbreaks, Viruses, Phylogeny, Poultry

## Abstract

In December of 2016, low pathogenic avian influenza (LPAI) caused by an H7N6 subtype was confirmed in a grow-out turkey farm located in Valparaiso Region, Chile. Depopulation of exposed animals, zoning, animal movement control and active surveillance were implemented to contain the outbreak. Two weeks later, a second turkey grow-out farm located 70 km north of the first site was also infected by H7N6 LPAI, which subsequently spilled over to one backyard poultry flock. The virus involved in the outbreak shared a close genetic relationship with viruses collected from Chilean aquatic birds’ viruses. The A/turkey/Chile/2017(H7N6) LPAI virus belonged to a native South American lineage. Based on the H7 and most of the internal genes phylogenies, these viruses were also closely related to the viruses that caused a highly pathogenic avian influenza outbreak in Chile in 2002. Results from this study help understand the regional dynamics of influenza outbreaks, highlighting the importance of local native viruses circulating in the natural reservoir hosts.

## INTRODUCTION

Outbreaks caused by avian influenza viruses (AIV) have major impacts in the poultry industry. Each year, events of highly pathogenic avian influenza (HPAI), and low pathogenic avian influenza (LPAI) are reported by different countries regardless of their overall animal health status and strategies to prevent the introduction of foreign animal diseases. These outbreaks cause high economic losses to the poultry industry and governments, due to elimination of all exposed animals, deployment of resources for surveillance and disease control strategies, as well as international trade restrictions (Djunaidi & Djunaidi, 2007; Otte, Hinrichs, Rushton, D, & D, 2008). Disease prevention is complex because many wild aquatic birds are a natural reservoir of influenza A viruses (Vandegrift, Sokolow, Daszak, & Kilpatrick, 2010).

The H5 and H7 subtypes are known for their potential to mutate to HPAI, typically after infecting poultry, resulting in severe disease and high mortality (Alexander, 2000). Among the outbreaks caused by H7, special attention has been given to two HPAI events. The first one was caused by an H7N9 virus that emerged in China in 2013. This virus is highly poultry adapted and spread widely in China, mutating to a highly pathogenic form during late 2016. The H7N9 virus that caused this outbreak is particularly concerning because, besides causing severe disease in poultry, it has also been responsible for more than 1,600 cases of illness in humans (CDC, 2018; Jiao et al., 2018; Ke et al., 2017). The second event also, caused by an H7N9 subtype was reported in March 2017 in the United States. This outbreak was caused by a North American wild bird lineage virus, which was genetically unrelated and epidemiologically different to the Asian H7N9 virus. Multiple LPAI introductions of this virus from wild birds into poultry were inferred based in epidemiological and molecular investigations. Mutation into HPAI occurred one time and resulted in the culling of more than 200,000 animals (Lee, Torchetti, Killian, Berhane, & Swayne, 2017). This event was contained and concluded six months after initial detection (OIE, 2018a).

The genetic diversity of AIV in South America has not been described in detail compared to that in North America. Regional and local dynamics of AIV have been associated to local South American and North American lineages and their transmission through migration routes that connect North and South America, especially the Pacific, Central and Mississippi Flyways. However, the information of subtypes and lineages circulating is not uniform across the continent, and throughout different time periods. Of the previously reported AIVs in Latin America, 43.7% belong to migratory birds, 28.1% to local wild birds, and 28.1% to poultry (Afanador-Villamizar, Gomez-Romero, Diaz, & Ruiz-Saenz, 2017). Additionally, several influenza subtypes have been detected in Chilean wild birds, including H5 and H7 subtypes, highlighting the risk for poultry production (Bravo-Vasquez et al., 2016; Jimenez-Bluhm et al., 2018; Mathieu et al., 2015).

Chilean poultry has been affected by HPAI only once, in 2002. The outbreak was caused by an HPAI A/chicken/Chile/2002(H7N3) virus that resulted in economic losses of over 31 million USD and ~635,000 animals dead or culled (Afanador-Villamizar et al., 2017; Max, Herrera, Moreira, & Rojas, 2007; Spackman, McCracken, Winker, & Swayne, 2006). Other AIV impacting poultry in Chile include an outbreak in turkeys in Valparaiso region in 2009, which was caused by the H1N1 pandemic (pH1N1) human virus. This event was evidenced by decreased egg production and shell quality, and was reported only two months after the first pH1N1 detection in humans in Chile (Mathieu et al., 2010). In 2011, LPAI H4N8 was found by serologic surveillance in a grow-out turkey farm, also in the region of Valparaiso. The presence of this AIV was confirmed by isolation in embryonated eggs. There was no evident clinical disease at the farm and rapid detection prevented further spread of the virus (SAG, 2011).

In this study, we describe the epidemiological investigations of an LPAI H7N6 outbreak in Chilean poultry in late 2016 and early 2017. We characterized the viruses using whole genome sequencing and reconstructed their phylogeny using Bayesian time divergence estimation, to infer their genetic relation with North and South American AIVs. These results highlight the importance of characterization and molecular surveillance of local South American AIV strains to understand the viruses that pose a major risk to domestic poultry.

## METHODS

### Description of the outbreak

On December 26^th^ of 2016 a grow-out turkey house (Farm A) in the region of Valparaiso reported clinical respiratory disease. On the following day, respiratory signs increased in severity. Noticeable, this farm had undergone serological (ELISA) AIV testing (n = 60) on December 20^th^, yielding negative results. On December 29^th^ and following the diagnosis of clinical disease, the poultry company notified positive AIV ELISA test results to the Chilean Agricultural and Livestock Service (Servicio Agricola Ganadero; SAG).

On December 30^th^, SAG officially confirmed the presence of AIV by real time RT-PCR (AI matrix gene). Subsequently, H5 and H7 subtype real time RT-PCR assays were performed, confirming an H7 subtype. The samples were initially sent to a private laboratory (Macrogen, Inc) for partial sequencing of the HA cleavage site. The amino acid sequence was consistent with a LPAI virus (corresponding to the translated amino acid site: NVPEKPRTR/GLF). Characterization of the neuraminidase gene subtype was carried out by neuraminidase-inhibition assay at SAG, revealing a N6 subtype. Further characterization of the virus included sequencing and *in vivo* testing to confirm the pathotype, performed at NVSL and SAG.

Once the LPAI H7N6 outbreak was officially confirmed, immediate control measures were established. A control area consisting of two zones surrounding the infected premises were defined: (1) an infected zone within a 3 km radius of the infected premises where animal movement was restricted, and (2) a buffer zone within 7 km radius of the infected premises, where increased biosecurity and surveillance for avian influenza were carried out (USDA, 2015). Additionally, all breeders and grow-out farms belonging to the affected company throughout the country were tested for influenza (including the ones outside the buffer zone). Surveillance consisted in blood sample collection and testing by agar gel immunodiffusion (AGID) assay performed at the SAG official diagnostic laboratory. A total of 327,000 exposed animals in Farm A were culled.

By January 2^nd^, clinical respiratory disease was present in flocks in five out of 8 houses within Farm A. The animals presented a variety of gross lesions including bursitis, catarrhal to mucopurulent tracheitis, caseous airsacculitis, polyserositis, pericarditis/hydropericardium, congestion and pulmonary oedema, mucopurulent to caseous pneumonia, localized subcutaneous emphysema, mild splenomegaly, and pancreatitis. Perihepatitis was only found in one house in the Farm A.

At the national level, recent records with productivity data from all commercial farms were reviewed, focusing in mortality, meat production, and egg production curves. The objective was to determine any abnormalities and potential undetected cases using syndromic surveillance. The parameters from all commercial farms yielded within-expected production values.

On 17^th^ January of 2017, 70 km north from the first affected farm (Valparaiso), a male turkey grow-out farm (Farm B) belonging to the same company, was visited by a veterinarian due to increased respiratory disease signs. SAG confirmed the presence of AIV and culled 35,572 exposed animals in the farm, which was subsequently cleaned and disinfected. Infected and buffer zones were put in place around the infected premises. Disposal of all carcasses was done by burial. In Farm B clinical gross lesions observed consisted of caseous sinusitis, catarrhal tracheitis, petechiae and ecchymosis focused on epicardium and coronary fat, fibrin-purulent pericarditis, purulent airsacculitis, pulmonary congestion/mucopurulent to caseous pneumonia, pleuritis, mild splenomegaly.

Epidemiological investigations concluded that the initial entry route of the virus into Farm A was through a breach in biosecurity which resulted in transmission from wild birds through faeces or contaminated water into one of the grow-out sites. Lateral transmission between Farm A and Farm B likely occurred due to an unreported breach in biosecurity by personnel, supplies or vehicles that were shared between Farm A and Farm B.

In January 28^th^ a backyard poultry farm within a household located in the defined buffer zone of Farm B tested positive to AIV by AGID but was negative to PCR testing. This farm had direct links through personnel with Farm B. All poultry in the household were culled.

Thereafter, all testing conducted in commercial farm inside and outside the control zones were negative to AIV. On June 9^th^, three months after finalizing stamping out of exposed and infected animals, Chile regained the OIE free status (OIE, 2017).

### Viral sequencing and phylogenetic analysis

Complete genome sequencing attempts were made from oral and tracheal swab pools at the United States Department of Agriculture, Animal and Plant Health Inspection Service, National Veterinary Services Laboratories, Ames, Iowa.

Briefly, viral RNA was extracted from samples using the MagMAX Viral RNA Isolation Kit (Ambion/ThermoFisher Scientific). Complementary DNA was synthesized by reverse transcription reaction using SuperScript III (Invitrogen/ThermoFisher Scientific). All eight segments were amplified by PCR and complete genome sequencing was conducted using the Illumina Miseq system. The Nextera XT DNA Sample Preparation Kit (Illumina) was used to generate multiplexed paired-end sequencing libraries, according to the manufacturer’s instructions. The dsDNA was fragmented and tagged with adapters by Nextera XT transposase and 12-cycle PCR amplification. Fragments were purified on Agencourt AMpure XP beads (Beckman Coulter). The barcoded multiplexed library sequencing was performed using the 500 cycle MiSeq Reagent Kit v2 (Illumina). De Novo and directed assembly of genome sequences were carried out using the SeqMan NGen v.4 program. Viral sequences were submitted to GenBank (accession numbers MK424141-MK424219).

Phylogenetic analyses were performed separately for each segment. The closest references viruses were obtained by a BLAST (https://blast.ncbi.nlm.nih.gov/Blast.cgi) search and the 100 closest hits were included to reconstruct the phylogenetic tree. Sequences obtained for each segment and the corresponding reference sequences were aligned MUSCLE (Edgar, 2004). Redundant reference sequence discarded (identical or near identical sequences with similar collection dates and locations). Bayesian time divergence estimation using the HKY+G[4] nucleotide substitution model and an uncorrelated relaxed clock was performed for all segments. Coalescent Bayesian Skyline tree prior was used to reconstruct the phylogeny of H7 (segment 4) to allow for population size changes over time. For N6 (segment 6) exponential growth and Bayesian skyline coalescent tree priors were compared using path sampling and stepping-stone sampling marginal likelihood estimation (Baele, Li, Drummond, Suchard, & Lemey, 2013). For the remaining segments 1, 2, 3, 5, 7 and 8 (coding for PB2, PB1, PA, NP, M and NS respectively) a coalescent exponential growth tree prior was initially run. If the 95% high posterior density (95% HPD) of the growth parameter included 0 (thus, meaning that the population size is not growing in time) a coalescent constant population tree prior was run and used for the final phylogeny, whereas if the parameter did not include 0, the original run with the exponential growth tree prior used was used. The analyses were run in BEAST 1.8.4 (Drummond, Suchard, Xie, & Rambaut, 2012). A total of 500 000 000 iterations were run, sampling every 50 000 trees using Cipres computational resources (Miller, Pfeiffer, & Schwartz, 2010). Traces of the parameters were checked for convergence and for effective sample size (ESS) > 200. The maximum clade credibility (MCC) tree was annotated, burning the first 10% sampled trees, using TreeAnnotator and visualized using Figtree (http://tree.bio.ed.ac.uk/software/figtree/). Amino acid sequences were compared to the closest references to identify relevant mutations using MEGA7 (Kumar, Stecher, & Tamura, 2016).

## RESULTS

Phylogenies were estimated using coalescent constant population tree priors for PB2, PA, NP, M and NS coding segments. Bayesian skyline tree prior was used for reconstructing HA and NA phylogenies, while exponential growth was used to estimate the phylogeny of PB1 (Table 1). The estimated nucleotide substitution rate per site per year for each coding segment ranged between 1.96x10^-3^ (95%HPD 1.52x10^-3^ - 2.44x10^-3^) and 3.98x10^-3^ (95%HPD 2.83x10^-3^- 5.26x10^-3^), which were the lowest and highest rates estimated for segment 7 (M) and 4 (HA), respectively.

**Table 1.**
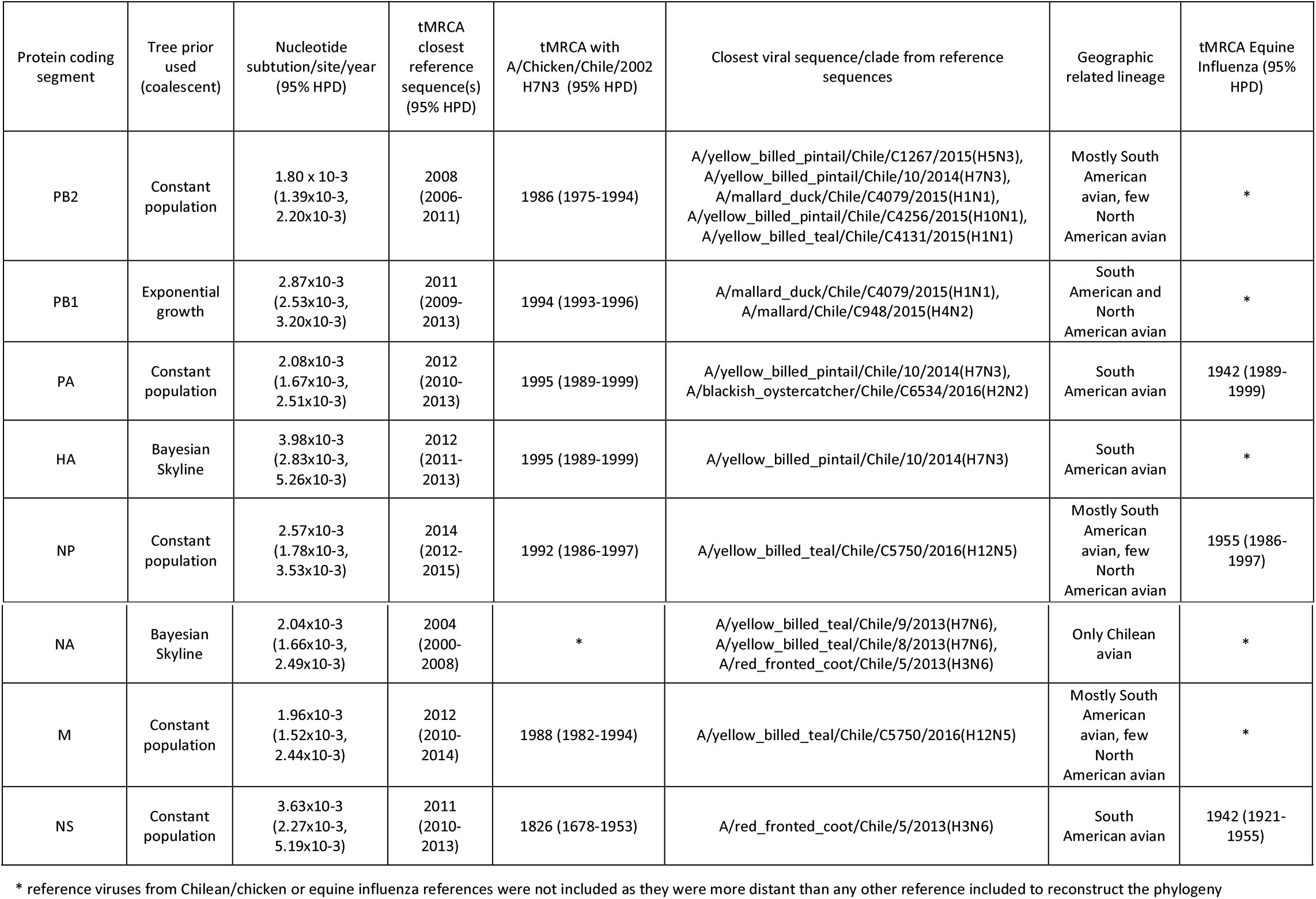
Results of Bayesian time estimation analyses for each protein-coding segment. Tree prior used for Bayesian time divergence estimation analysis for each segment alignment is shown. The estimated nucleotide substitution rate per site per year for each coding segment was highest for HA and lowest for M segment. The time to the most recent common ancestor (tMRCA) with the closest reference virus, with A/chicken/Chile/2002(H7N3) and with equine influenza virus are indicated for related viral segments, as well as the characteristics of the lineage to which they belong.

### HA coding segment

All viruses from the 2017 Chilean turkey LPAI outbreak viruses were included in a monophyletic clade, which shared the closest ancestor with 2 viruses (A/yellow_billed_pintail/Chile/10/2014(H7N3), and A/yellow_billed_teal/Chile/9/2013(H7N6)) collected from wild local water birds. The tMRCA shared by the closest Chilean wild bird virus and the A/turkey/Chile/2002(H7N6) was estimated at 2012 (95% HPD 2011-2013) (Figure 1). The A/turkey/Chile/2017(H7N6) LPAI viruses were also related to AIV found in wild aquatic birds in Chile and Bolivia, and with the 2002 HPAI Chilean outbreak with a tMRCA estimated at 1995 (95% HPD 1989-1999). Based on the HA phylogeny, these South American viruses were grouped in a unique cluster. The remaining closest reference, belonging to a North American avian lineage, that recently caused an HPAI outbreak in the United States, shared an earlier common ancestor estimated in 1944 (95% HPD 1913-1969). Specific amino acid changes fixed in the LPAI A/turkey/Chile/2017(H7N6) compared to the closest references were in HA amino acid positions V42I, V96I, I191V, P524S, and a fixed mutation in Farm B sequences R316K (Relative to HPAI Chicken/Chile2002 which has a ten-amino acid insertion in positions 338-347; Table 2). Three of these specific amino acid changes are located in antigenic sites C, D and E (Supplementary file 1), which are not related to additional glycosylation sites.

**Figure 1.**
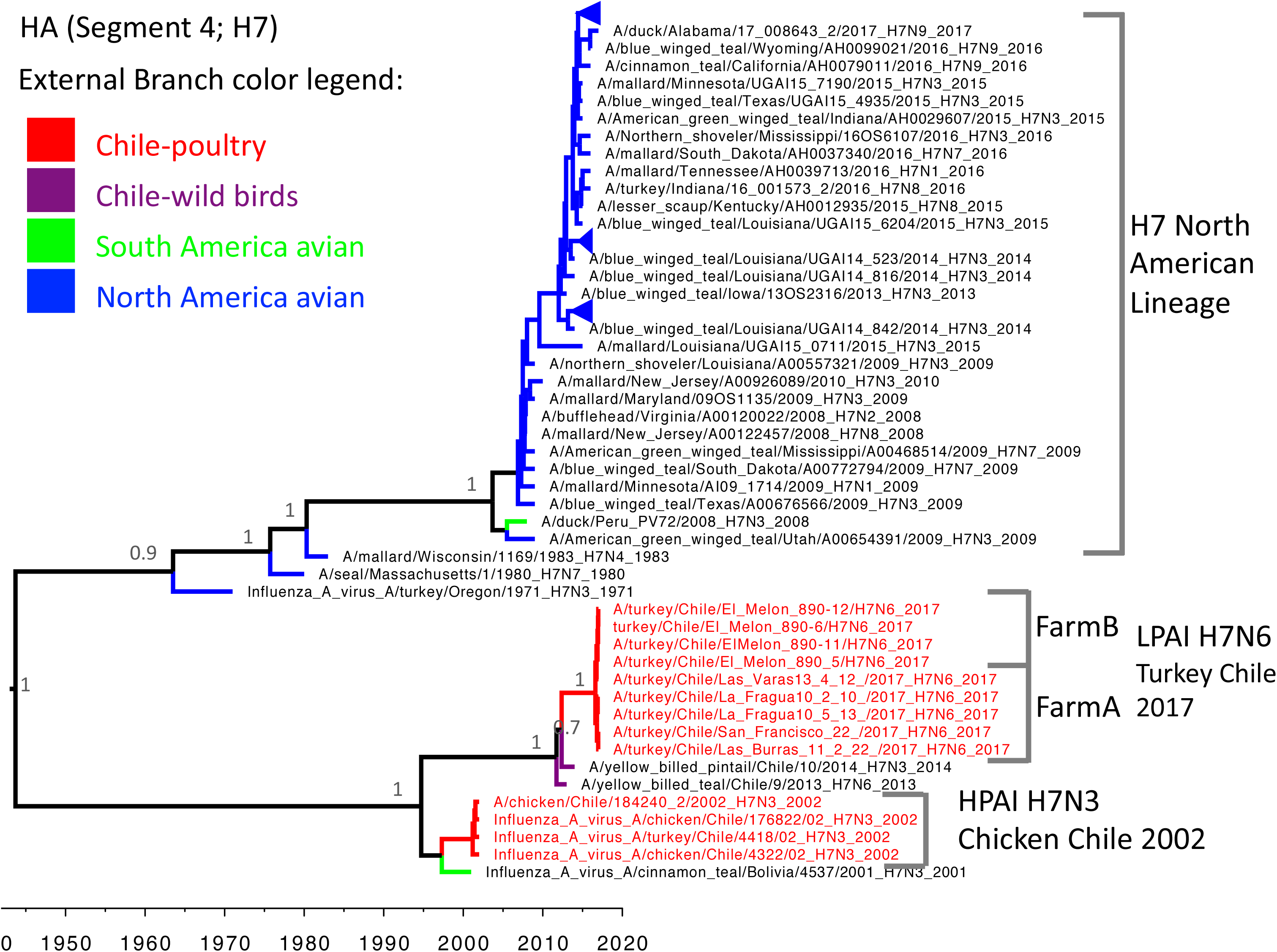
Maximum clade credibility tree of HA segment. The South American lineage shared a common ancestor in 1944 (95% HPD 1913-1969) with the closest reference viruses (North American Lineage). The viruses that caused the H7N6 LPAI outbreak in 2017 in Chile were genetically related to the viruses that caused an HPAI outbreak in 2002 in Chile (A/chicken/Chile/2002(H7N3)).

**Table 2.**
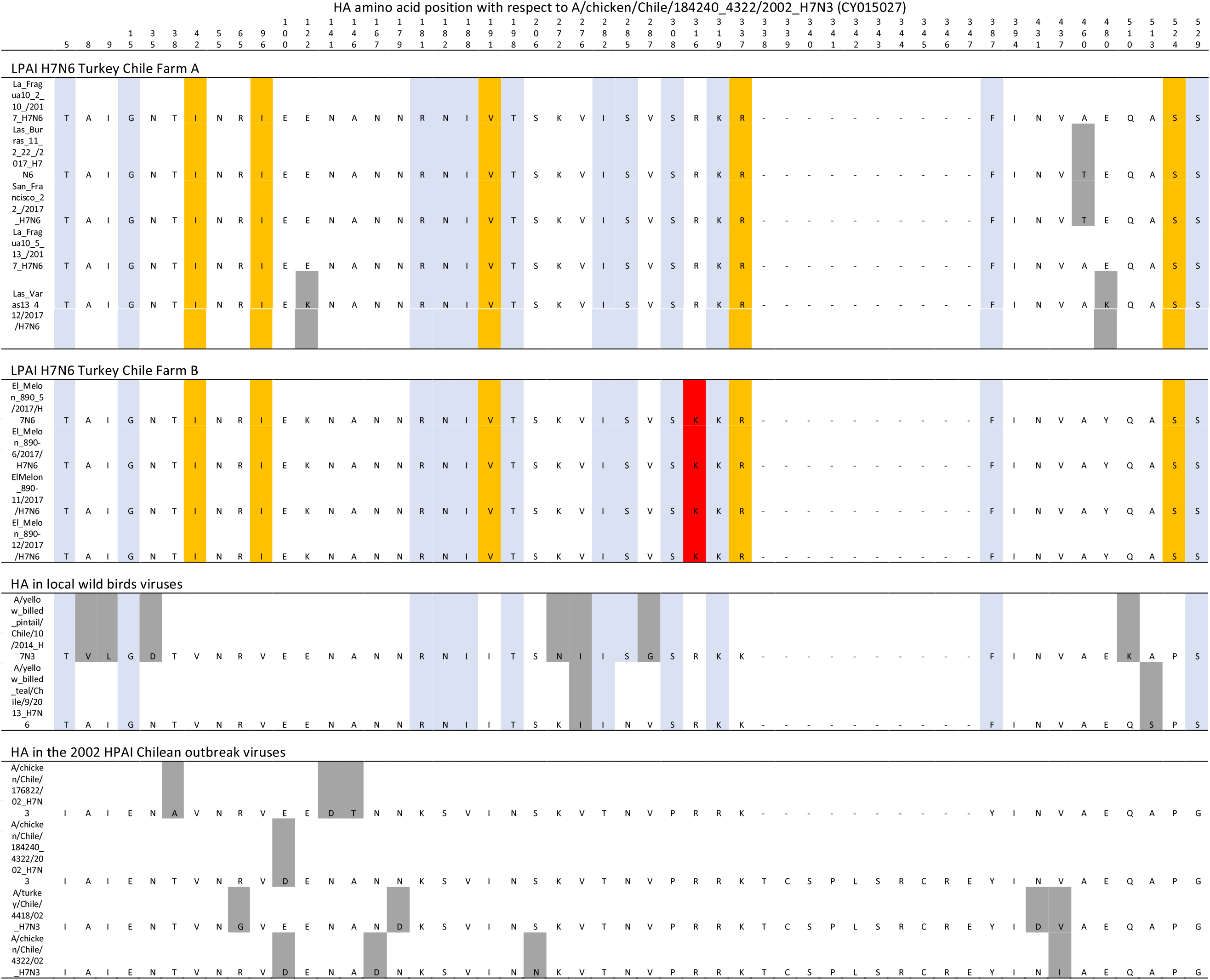
Amino acid difference in HA coding segment between the A/turkey/Chile/2017(H7N6), the closest viruses collected from wild birds, and viruses from the previous Chicken/Chile/2002 H7N3 outbreak. Changes fixed in viruses collected after the 2002 outbreak are coloured in blue, whereas changes fixed only in A/turkey/Chile/2017(H7N6) LPAI are highlighted in orange. Non-fixed mutations are highlighted in grey. One nucleotide substitution was fixed in Farm B R316K (highlighted in red).

### NA coding segment

Similar to HA coding segments, NA sequences from the A/turkey/Chile/2017(H7N6) LPAI outbreak are genetically related to a distinct group that includes viruses collected from Chilean wild birds in 2013 (H7 and H3 subtypes being the closest ones) with a tMRCA estimated at 2004 (95% HPD 2000-2008) (Figure 2). This group is more genetically distant to viruses from North American avian compared to the HA and most internal genes’ phylogenies, having a tMRCA estimated at 1869 (95% HPD 1823-1911). Sequences from A/turkey/Chile/2017(H7N6) LPAI had three amino acid substitutions fixed compared to the closest references (amino acid positions T42P, V333A and N359S). We included only one sequence per sample to depict the phylogeny, however, a mixed population of NA sequences with and without stalk deletions (ranging between 23-27 amino acid long) were found in (pooled) samples collected from Farm A. All four samples from Farm B had sequences with the same 26 amino acid nucleotide stalk deletion between amino acid position 32-57. This deletion was not associated with additional glycosylation sites in HA.

**Figure 2.**
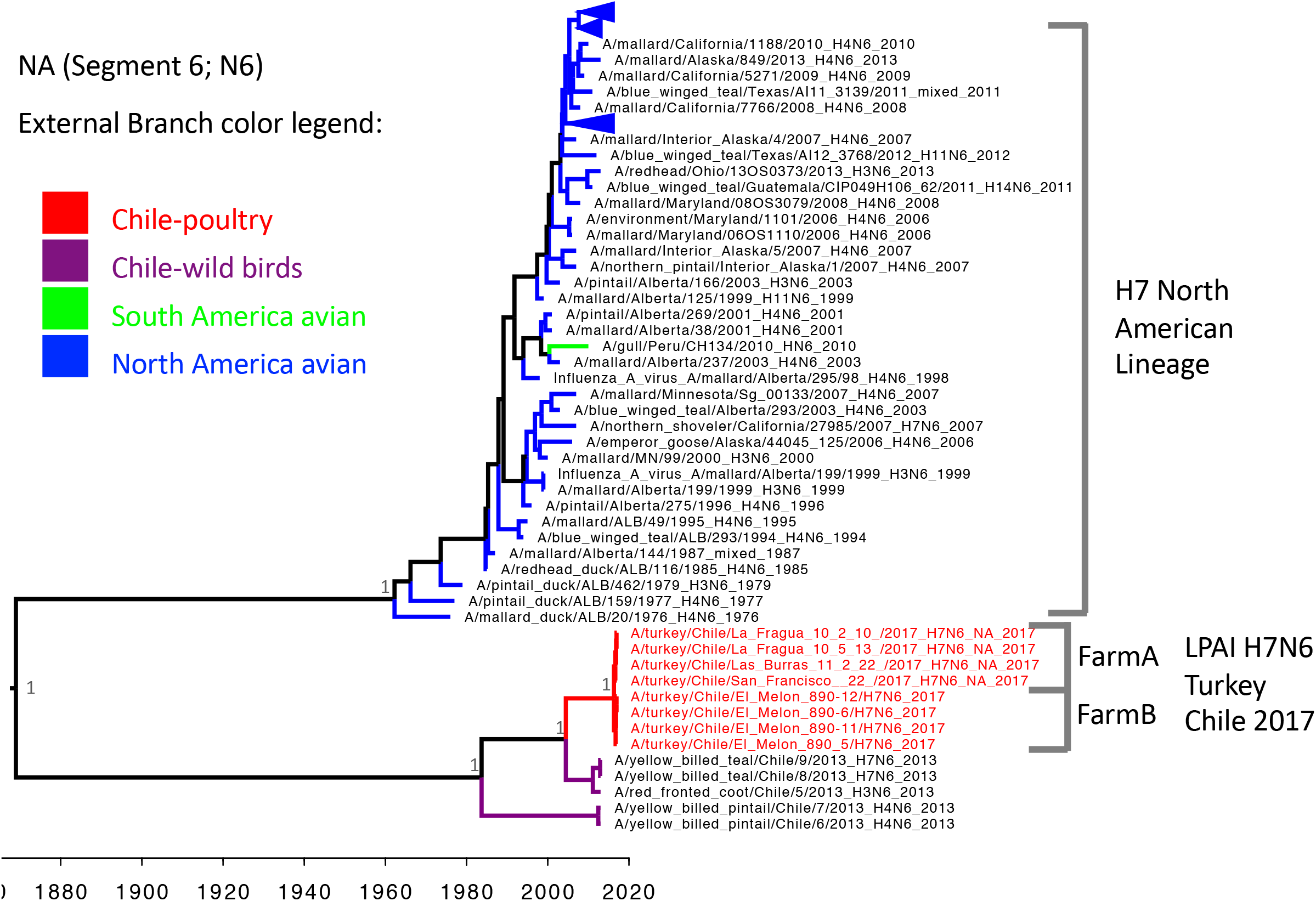
Maximum clade credibility tree of NA segment. A/turkey/Chile/2017(H7N6) LPAI viruses are grouped in a unique cluster unrelated to other references and sharing a common ancestor in 1869 (95% HPD 1823-1911) with the closest North American lineage.

### Phylogeny of internal segments PB2, PB1, PA, NP, M and NS

For the reconstructed phylogeny of internal segments, the closest reference sequences to the A/turkey/Chile/2017(H7N6) LPAI viruses were viruses collected from Chilean aquatic birds (Figure 3). The closest relationships for each segment of A/turkey/Chile/2017(H7N6) LPAI were with different viruses from a variety of HA and NA types (Table 1). PB1, PA, HA and NP internal genes coding segments of A/turkey/Chile/2017(H7N6) LPAI shared a similar tMRCA with A/chicken/Chile/2002(H7N3) HPAI estimated between 1992-1995. In terms of regional lineages, the A/turkey/Chile/2017(H7N6) LPAI segments PB2, NP and M were grouped with avian viruses sampled mostly from South America and few from North America, whereas segments PA, HA, and NS clustered with sequences that were exclusively South American. The PA, NP and NS lineages to which A/turkey/Chile/2017(H7N6) LPAI belong, were related to equine viruses from as a result from a potential reassortment event that occurred sometime between 1942-1955 (Table 1). Interestingly, one of the M segments from the A/chicken/Chile/2002(H7N3) outbreak was distinct from other viruses collected during that epidemic (Figure 3).

**Figure 3.**
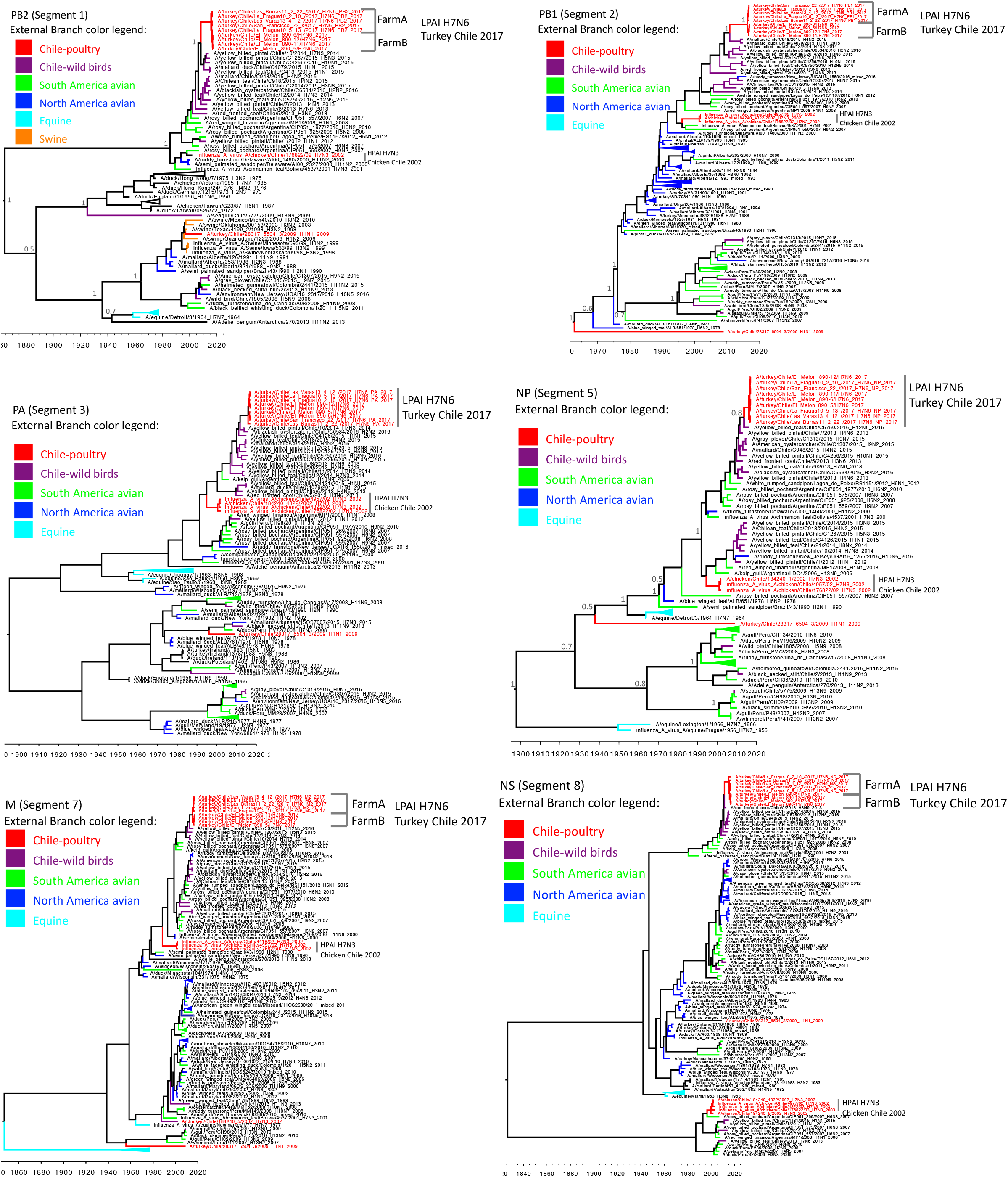
Maximum clade credibility tree of segments PB2, PB1, PA, NP, M and NS. All segments are closely related to viruses previously collected from Chilean wild birds. All Segments, except for NS, are related to the previous Chilean 2002 HPAI outbreak.

Nucleotide mutations fixed in the A/turkey/Chile/2017(H7N6) LPAI viruses compared to close reference from local aquatic birds were found in PB1 (V344I, V444I) and T683I, in M2 (K18R, V27I) and in NP (V193I, T196I, M221V) as well as a Farm B specific fixed mutation in PB1 (R101G, N573H) and NP (M123I).

None of the segments were related to the pH1N1 viruses that caused an outbreak in turkeys during 2010, and which HA and NA were related to the 2009 pandemic virus.

## DISCUSSION

In Chile, an outbreak of H7N6 LPAI (A/turkey/Chile/2017(H7N6)) was detected shortly after initial signs of respiratory disease were reported in a commercial turkey grow-out farm in December of 2016. Epidemiological investigations and phylogenetic analyses of viral sequence data were used to understand the source of the outbreak and characterize the lineage and the genetic relationships with reference viruses. All segments from the A/turkey/Chile/2017(H7N6) LPAI virus were related to viruses collected in previous years from wild birds in Chile. The A/turkey/Chile/2017(H7N6) LPAI virus grouped within the same lineage as the Chicken/Chile/2002(H7N3) HPAI, based on the phylogeny of HA and most genetic segments coding for internal proteins. The A/turkey/Chile/2017(H7N6) LPAI virus belongs to a native South American lineage, and are not closely related to other recent viruses reported from poultry in the Americas such as the subtype H7N9 that had caused outbreaks during 2017 in commercial chicken farms in Tennessee and Alabama in the United States (Lee et al., 2017), and the H7N3 that has caused sporadic HPAI outbreaks in Mexico since 2015 (OIE, 2018b). This South American lineage is also unrelated to the poultry adapted Asian H7N9 virus (CDC, 2018).

Control of AIV in poultry is complex and challenging. High biosecurity measures must be accomplished to prevent viruses that continuously circulate in wild birds from entering poultry operations. A wide diversity of influenza A viruses exist among water birds, which may persist in geographical areas where commercial and backyard poultry farms are located. A large number of wetlands throughout the extensive Chilean coast, provides an ecosystem where migratory and local wild water birds (including the main avian influenza reservoirs) co-exist (Jimenez-Bluhm et al., 2018). Small backyard production systems are also constantly exposed to transmission from wild birds and are, likely, sporadically infected (Bravo-Vasquez et al., 2016), representing another potential source of AIV for commercial farms. Additionally, transmission from human or other species also pose a risk for influenza outbreaks in poultry, such as the pH1N1 introduced from human into a commercial turkey farm in Chile in 2009 (Mathieu et al., 2010).

A close ancestral relationship between viruses that caused the A/chicken/Chile/2002(H7N3) HPAI and the A/turkey/Chile/2017(H7N6) LPAI revealed the importance of this specific South American lineage virus as a risk for commercial poultry farms. Aquatic bird reservoirs have maintained this virus throughout the years, posing a risk to poultry farms that was not evidenced until the recent 2017 outbreak. Considering this H7N6 event, as well as the previous H7N3 in Chile, this local viral lineage has been the main source of AIV poultry outbreaks. However, the risk of introduction of other lineages into Chile should not be underestimated as there is a wide variety of local and North American lineages found in wild water birds.

Previous studies have supported that A/chicken/Chile/2002(H7N3), as well as other American AIV H7 subtype strains, need a lower infectious dose to cause clinical disease in turkey, compared to that needed to affect chicken (Spackman et al., 2010; Spackman et al., 2006). It is relevant to design targeted surveillance, as turkey may act as sentinels for emerging and more pathogenic AIV strains. In this particular event, clinical disease caused by LPAI in two turkey farms was rapidly detected and preventive measures were timely put in place to contain viral spread. Some viruses in Farm A and all viruses from Farm B had already acquired amino acid stalk deletion in its NA segment. These genetic changes have been associated to an adaptation to the domestic host and increased virulence and transmission (Campitelli et al., 2004; Li, Zu Dohna, Cardona, Miller, & Carpenter, 2011; Sorrell, Song, Pena, & Perez, 2010). Therefore, this virus was rapidly adapting with had a potential to further spread into other farms, if it had not been detected on time.

The genetic changes occurred in the viral genome within a short time window (less than a month) allowed to correctly group Farm A and B clusters in the phylogeny reconstructed for segments HA, NA PB1 and M. However, the reconstruction of the genetic similarity of the remaining segments did not allow to discern between Farm A and B. This is information is relevant to understand the level of resolution to reconstruct inter farm transmission networks in based on phylogenies.

Because of the rapid diagnostic and intervention, the LPAI outbreak that affected Chilean turkeys in 2017 was contained and limited to only 2 premises of the same company. Phylogenetic analyses revealed that the outbreak was caused by a South American wild bird lineage virus, and unrelated to the 2017 North American H7N9 outbreak in the United States in March. The lack of systematic molecular AIV surveillance in South American countries highlights the need to coordinate regional efforts to better understand the genetic diversity and viral dynamics in the region, and to be prepared for future outbreaks in domestic species (Afanador-Villamizar et al., 2017). This report contributes to understanding this particular clinical LPAI event in turkey and the role of local South American lineages circulating in the wild reservoirs as a major source of strains that are pathogenic to domestic poultry.

## ACKNOWLEDGEMENTS

We thank all personnel from the Virology Unit and Betty Yangari from Biotechnology Unit at the Lo Aguirre Agricultural and Livestock Service Laboratory. B. Brito was funded by the University of Technology Sydney under the Chancellor’s Research Fellowship Program. V. Neira was funded by Programa Fondecyt de Iniciación N° 11170877 and to Programa de Investigación Asociativa from the Comisión Nacional de Investigación Científica y Tecnológica, project CONICYT-PIA Anillo ACT 1408.

---

**Supplementary file 1.** Antigenic sites of in HA sequences of reference viruses and viruses collected from poultry and wild birds in Chile.

